# Regulatory T Cell-like Response to SARS-CoV-2 in Jamaican Fruit Bats (*Artibeus jamaicensis*) Transduced with Human ACE2

**DOI:** 10.1101/2023.02.13.528205

**Authors:** Bradly Burke, Savannah M Rocha, Shijun Zhan, Miles Eckley, Clara Reasoner, Amin Addetia, Juliette Lewis, Anna Fagre, Phillida Charley, Juergen A Richt, Susan R Weiss, Ronald B Tjalkens, David Veesler, Tawfik Aboellail, Tony Schountz

**Affiliations:** Department of Microbiology, Immunology and Pathology, College of Veterinary Medicine, Colorado State University, Fort Collins, Colorado USA; Department of Environmental and Radiological Health Sciences, College of Veterinary Medicine, Colorado State University, Fort Collins, Colorado USA; Department of Biochemistry, University of Washington, Seattle, WA 98195, USA; Diagnostic Medicine/Pathobiology, Center of Excellence for Emerging and Zoonotic Animal Diseases (CEEZAD), College of Veterinary Medicine, Manhattan, KS 66503 USA; Department of Microbiology, Perelman School of Medicine, University of Pennsylvania, Philadelphia, PA 19104 USA

## Abstract

Insectivorous Old World horseshoe bats (*Rhinolophus* spp.) are the likely source of the ancestral SARS-CoV-2 prior to its spillover into humans and causing the COVID-19 pandemic. Natural coronavirus infections of bats appear to be principally confined to the intestines, suggesting fecal-oral transmission; however, little is known about the biology of SARS-related coronaviruses in bats. Previous experimental challenges of Egyptian fruit bats (*Rousettus aegyptiacus*) resulted in limited infection restricted to the respiratory tract, whereas insectivorous North American big brown bats (*Eptesicus fuscus*) showed no evidence of infection. In the present study, we challenged Jamaican fruit bats (*Artibeus jamaicensis*) with SARS-CoV-2 to determine their susceptibility. Infection was confined to the intestine for only a few days with prominent viral nucleocapsid antigen in epithelial cells, and mononuclear cells of the lamina propria and Peyer’s patches, but with no evidence of infection of other tissues; none of the bats showed visible signs of disease or seroconverted. Expression levels of ACE2 were low in the lungs, which may account for the lack of pulmonary infection. Bats were then intranasally inoculated with a replication-defective adenovirus encoding human ACE2 and 5 days later challenged with SARS-CoV-2. Viral antigen was prominent in lungs for up to 14 days, with loss of pulmonary cellularity during this time; however, the bats did not exhibit weight loss or visible signs of disease. From day 7, bats had low to moderate IgG antibody titers to spike protein by ELISA, and one bat on day 10 had low-titer neutralizing antibodies. CD4^+^ helper T cells became activated upon ex vivo recall stimulation with SARS-CoV-2 nucleocapsid peptide library and exhibited elevated mRNA expression of the regulatory T cell cytokines interleukin-10 and transforming growth factor-β, which may have limited inflammatory pathology. Collectively, these data show that Jamaican fruit bats are poorly susceptibility to SARS-CoV-2 but that expression of human ACE2 in their lungs leads to robust infection and an adaptive immune response with low-titer antibodies and a regulatory T cell-like response that may explain the lack of prominent inflammation in the lungs. This model will allow for insight of how SARS-CoV-2 infects bats and how bat innate and adaptive immune responses engage the virus without overt clinical disease.

**Author Summary:** Bats are reservoir hosts of many viruses that infect humans, yet little is known about how they host these viruses, principally because of a lack of relevant and susceptible bat experimental infection models. Although SARS-CoV-2 originated in bats, no robust infection models of bats have been established. We determined that Jamaican fruit bats are poorly susceptible to SARS-CoV-2; however, their lungs can be transduced with human ACE2, which renders them susceptible to SARS-CoV-2. Despite robust infection of the lungs and diminishment of pulmonary cellularity, the bats showed no overt signs of disease and cleared the infection after two weeks. Despite clearance of infection, only low-titer antibody responses occurred and only a single bat made neutralizing antibody. Assessment of the CD4^+^ helper T cell response showed that activated cells expressed the regulatory T cell cytokines IL-10 and TGFβ that may have tempered pulmonary inflammation.

## Introduction

The ancestral severe acute respiratory syndrome coronavirus 2 (SARS-CoV-2), in all likelihood, originated in insectivorous bats prior to spillover to humans through one or more intermediate bridge hosts in live animal markets in Wuhan, China (1–4). SARS-CoV-2 is a sarbecovirus (genus *Betacoronavirus*, subgenus *Sarbecovirus*) that uses angiotensin converting enzyme 2 (ACE2) as a cellular entry receptor in humans. Although hundreds of distinct sarbecovirus sequences have been detected in bats in Asia and Europe, principally in horseshoe bats (*Rhinolophus* spp.) (3, 5–11), not all can use human ACE2 as an entry receptor (12).

Field studies of natural coronavirus infections of bats show that infections are localized to the gastrointestinal tract with likely shedding through feces, suggesting maintenance of virus in natural bat populations is via fecal-oral transmission (1, 5–7). However, few studies have experimentally examined coronavirus infections in bats, and those that have relied exclusively on surrogate models. Experimental challenge determined that Jamaican fruit bats (*Artibeus jamaicensis*), one of the most abundant and largest bats in the Americas, are susceptible to the merbecovirus, Middle East respiratory syndrome coronavirus (MERS-CoV) (13). Viral antigen and RNA were detected in several organs up to 14 days post challenge, and viral RNA was detected in oral and rectal swabs up to 9 days post challenge. Inoculation of Jamaican fruit bat primary kidney cells (Ajk cells) led to robust virus replication and cytopathic effect, with destruction of the Ajk cell monolayer. Despite clear evidence of infection, no conspicuous disease occurred, only one bat seroconverted, and they remained healthy. MERS-CoV likely emerged from insectivorous bats in Africa (14–16) that used dromedary camels (*Camelus dromedarius*), which then became a secondary reservoir host species (17), as a bridge to humans.

Considering the wide species tropism of SARS-CoV-2, which includes mustelids, cervids, felines, canines and cricetid rodents, including Syrian hamsters and North American deer mice (18–24), it might be expected that SARS-CoV-2 can readily infect a variety of bat species. A significant limitation for conducting meaningful studies on the biology of bat-borne viruses is the lack of relevant captive breeding colonies of specific pathogen-free (SPF) bats. A few established colonies of SPF fruit bats and nectarivorous bats are available for infectious disease studies (25–27); however, insectivorous bats are substantially more difficult to colonize because of the need for live insect species that are often not natural food sources of the bats. Thus, insectivorous bats are typically captured and used for one-off experimental infection studies. Despite these limitations, previous work using Egyptian fruit bats (*Rousettus aegyptiacus*), and insectivorous big brown (*Eptesicus fuscus*) and Brazilian/Mexican free-tailed (*Tadarida brasiliensis*) bats have examined susceptibility to SARS-CoV-2 infections (28–31). Although Egyptian fruit bats were moderately susceptible without disease, with detection of virus in the respiratory tract for several days and transmission to one other bat, big brown bats were not susceptible. There is conflicting evidence of susceptibility of Brazilian free-tailed bats; one study suggested they are not susceptible (30), whereas another suggested low susceptibility (31). Thus, it is clear host specificity of SARS-CoV-2 infection cannot be determined simply by taxonomic assignment of a given mammalian species.

We sought to determine the susceptibility of Jamaican fruit bats to SARS-CoV-2. Initial challenge of bats with SARS-CoV-2 WA1 isolate was confined to the small intestine for only 2 days before antigen could no longer be detected, bats held 21 days failed to seroconvert, and contact transmission did not occur. Viral antigen or RNA was not detected in other tissues, suggesting an abortive infection that is poorly adapted to Jamaican fruit bats, that infection was controlled by the innate immune response, or both. We were interested in determining whether ACE2 was a restriction factor for SARS-CoV-2 infection of Jamaican fruit bats; therefore, we intranasally inoculated bats with an adenovirus that expresses human ACE2 (32–34) and subsequent challenge of the bats with SARS-CoV-2 led to robust infection in the lungs for up to 14 days and low-titer seroconversion. Histopathology revealed moderate inflammation in the lungs, including neutrophil infiltration, lymphoblasts and macrophage syncytia, and loss of pulmonary cellularity. Bats appeared healthy and experienced no weight loss, suggesting only mild disease. Splenic CD4^+^ helper T cells were activated upon ex vivo stimulation with nucleocapsid peptide library and three of four bats expressed the regulatory T cell cytokines IL-10 and TGFβ but not inflammatory cytokines. Collectively, this study shows that Jamaican fruit bats are poorly susceptible to SARS-CoV-2; however, they are highly susceptible when a suitable ACE2 receptor is provided but which only leads to minimal pathology, and that a virus-specific adaptive immune response occurs that may be mediated by regulatory T cells. Importantly, this study provides the first example of a robust infection of bats with SARS-CoV-2, virus-specific bat T cell responses, and demonstrates the use of adenovirus vectors for in vivo expression of genes of interest in Jamaican fruit bat lungs.

## Results

### SARS-CoV-2 causes restricted infection of Jamaican fruit bats that is confined to the small intestine

Intranasal inoculation of Jamaican fruit bats with SARS-CoV-2 WA1 did not result in visible signs of disease nor did they lose weight. However, viral antigen was readily detected in intestinal sections on day 2 (Figure 1A) but not day 4 or thereafter. Antigen was particularly prominent in the periphery of Peyer’s patches that are typically populated with macrophages, mononuclear cells of the lamina propria, and intestinal crypts. No antigen was detected in intestinal sections of unchallenged bats (Figure 1B). Oral swabs, but not rectal swabs, from bats were positive for viral RNA (vRNA) on days 2 and 4 post challenge (Figure 1C), but negative thereafter. Despite clear evidence of antigen staining, both ELISA to recombinant nucleocapsid antigen and virus neutralization assay were negative for antibodies. To determine why virus was not detected in the lungs, we examined ACE2 gene expression levels and found that it was very low (Figure 1D), suggesting this may be a restriction factor for infection in the lungs.

**Figure 1.**
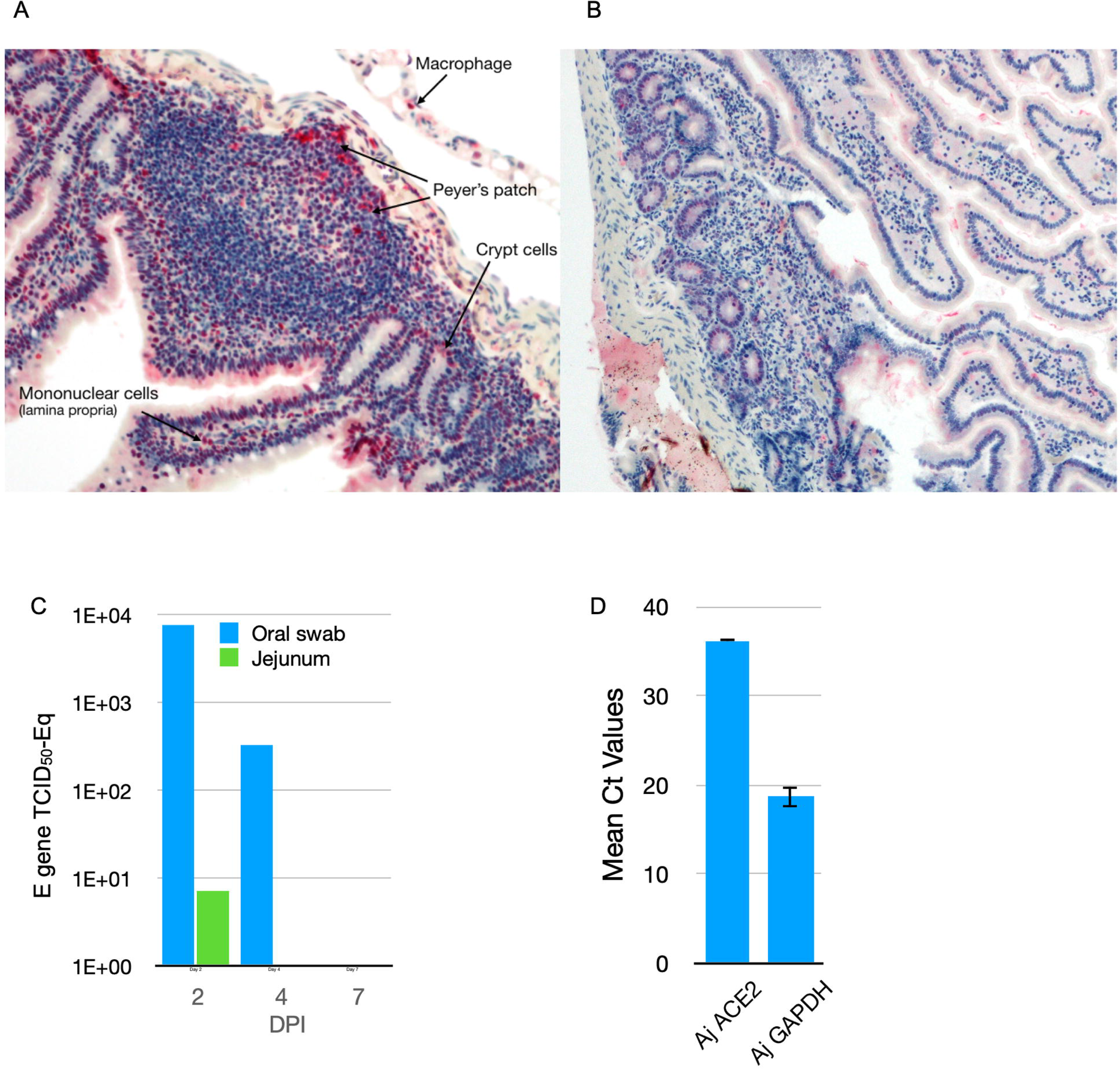
Immunohistochemistry of intestines of Jamaican fruit bats challenged with SARS-CoV-2. **A**. Two days after challenge, but not thereafter, antigen was detected in mononuclear cells of the lamina propria and periphery of Peyer’s patches where macrophages are typically found. Crypt cells also were infected. **B**. Unchallenged control bat intestine showed no antigen staining. **C**. SARS-CoV-2 E gene vRNA was detected only on days 2 and 4 in oral swabs, and only on day 1 in jejunum. **D**. Endogenous ACE2 expression in lungs of Jamaican fruit bat appears to be low and may account for the lack of detectable virus in lungs.

### Defective adenovirus encoding human ACE2 transduces Jamaican fruit bat cells

A replication-defective adenovirus serotype 5 that encodes human ACE2 (Ad5/hACE2) has been used to study SARS-CoV-1 and SARS-CoV-2 infections of nonsusceptible laboratory mice (32, 33). Examination of the mouse receptor for Ad5, the coxsackievirus and adenovirus receptor (CAR), revealed that it has 85% identity and 92% similarity to Jamaican fruit bat CAR, suggesting that Jamaican fruit bats may be susceptible to Ad5. To test this hypothesis, we inoculated primary Jamaican fruit bat kidney (Ajk) cells with Ad5/hACE that also encodes eGFP and examined the cells daily. By 24 hours, florescence was observed and it was maximal on days 2 through 6 when the study was terminated (Figure 2A). Ajk cells, transduced for 48 hours, and Vero E6 cells (positive control) were inoculated with 0.1 MOI of SARS-CoV-2 and supernatants were collected at 1 hour and daily for 4 days, followed by titration on Vero E6 cells. No increase in SARS-CoV-2 was detected from hACE2-transduced Ajk cells (Figure 2B), suggesting the cells were not permissive for SARS-CoV-2 replication.

**Figure 2.**
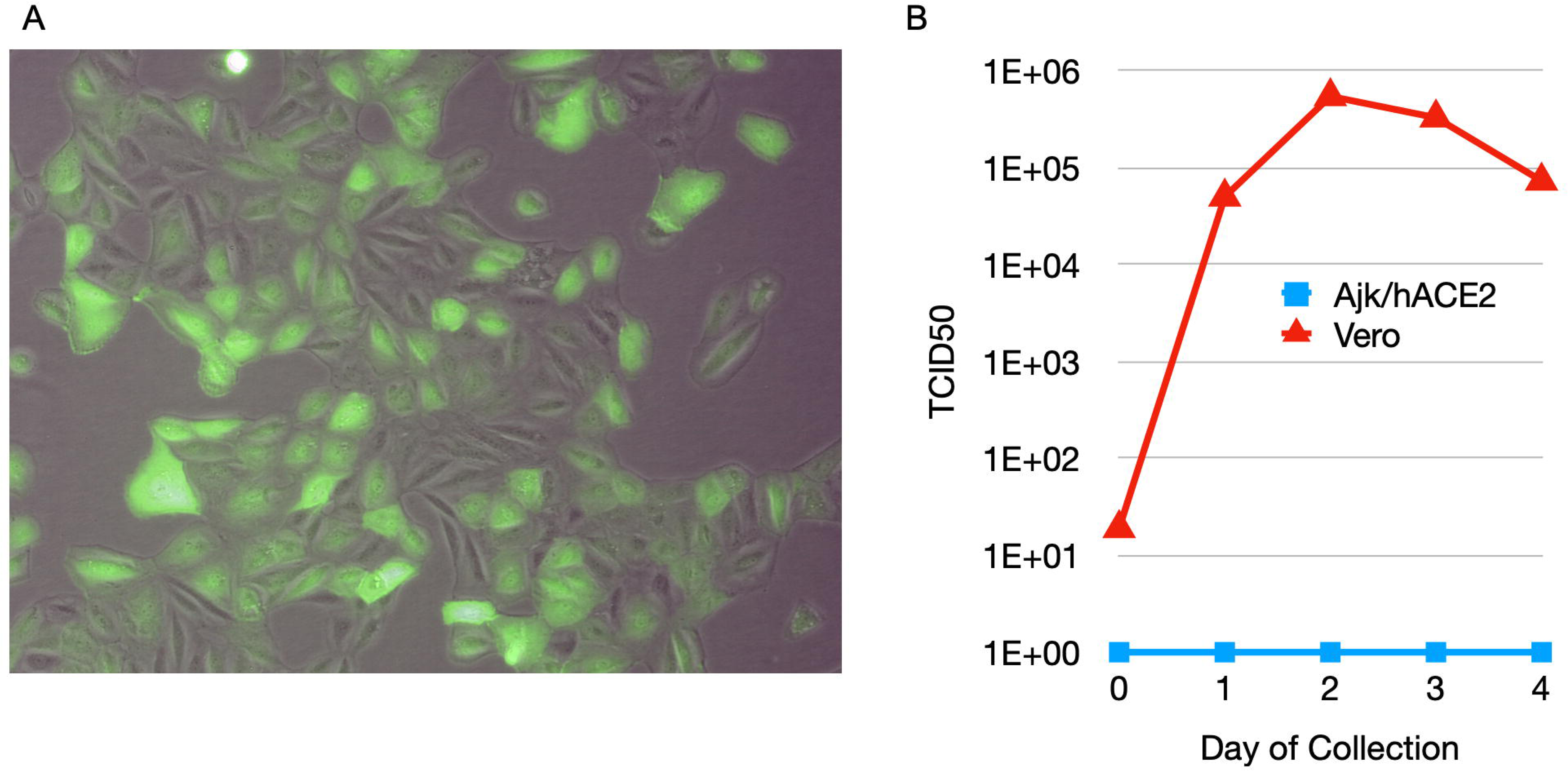
Adenovirus serotype 5 encoding human ACE2 (Ad5/hACE2) transduces Jamaican fruit bat cells in vitro. **A.** Transduction of Jamaican fruit bat primary kidney epithelial (Ajk) cells with Ad5/hACE2/eGFP. Jamaican fruit bat primary kidney cells were inoculated with 10 MOI of Ad5/hACE2/eGFP for 1 hour, washed and incubated in fresh medium for 4 days. Most, but not all, cells exhibited fluorescence, suggesting a heterogeneous population with some susceptible and some nonsusceptible cells. **B**. Ad5/ hACE2 transduced Jamaican fruit bat kidney cells are not productively infected with SARS-CoV-2. Ajk/ hACE2 cells (blue) and Vero cells (red) were inoculated with 0.1 MOI of SARS-CoV-2 WA1 for 1 hour, washed and cultured for 4 days. Supernatants were collected each day and titrated on Vero E6 cells to determine virus titers. Despite evidence of human ACE2 expression on Ajk cells, inoculation failed to produce infectious virus, suggesting the cells were not permissive.

### Bat lung cells transduced with human ACE2 are susceptible to SARS-CoV-2

To determine if lung cells of Jamaican fruit bats transduced with human ACE2 become susceptible to SARS-CoV-2, 21 bats were intranasally inoculated with 2.5×10^8^ pfu of Ad5/hACE2 and 5 days later they were challenged with SARS-CoV-2 (Figure 3A). Groups of bats (four on days 2, 4, 7; two on days 10, 14, 21) were serially euthanized. On day 2, lungs of all four euthanized bats had detectable viral RNA (vRNA); however, on days 4 and 7 only one bat from each group had vRNA, and none of the lungs from bats thereafter had vRNA (Figure 3B). Serum antibody to recombinant spike was detected by ELISA in one bat on day 4, and all bats euthanized thereafter, with peak response on day 10 (Figure 3C). Only one bat, euthanized on day 10, had weak, but detectable, neutralizing antibody using a D614G VSV pseudotype assay (Supplemental Figure 1); this bat also had the highest O.D. value on the ELISA (2.957). No weight loss occurred in the bats (Figure 3D).

**Figure 3.**
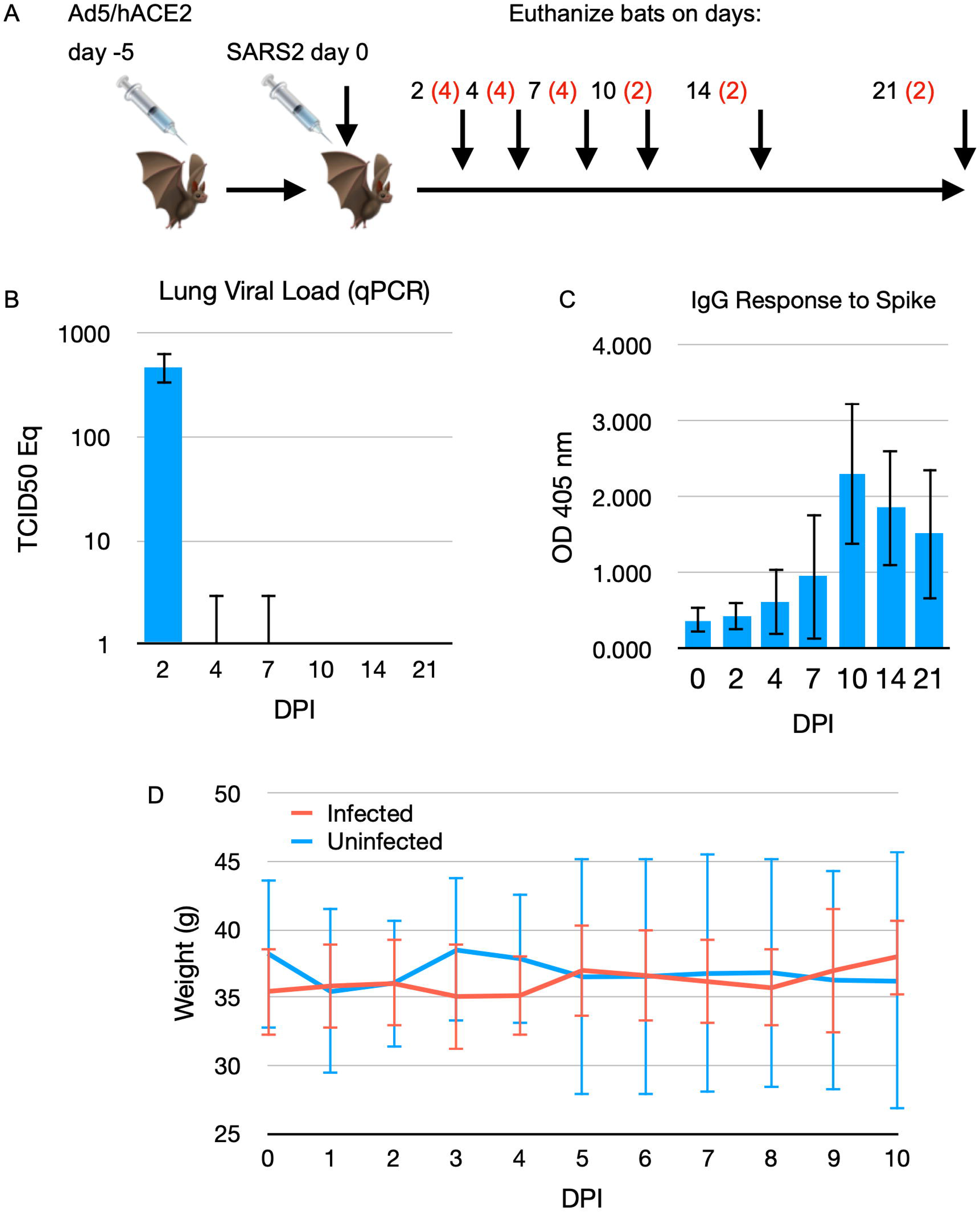
SARS-CoV-2 infection of Jamaican fruit bat lungs transduced with hACE2. **A.** Experimental design. Jamaican fruit bats were intranasally inoculated with Ad5/hACE2, then challenged with SARS-CoV-2 WA1. Groups of bats (n=red numerals) were euthanized at intervals and evaluated for infection. **B**. Viral RNA was detected in all bats on day 2 but only one bat on each of days 4 and 7. **C**. Serum antibody titers to recombinant spike antigen became elevated by day 4 in one bat, and all bats thereafter. **D**. Bats did not lose weight during the first ten days of the study, nor did they show other signs of disease.

### SARS-CoV-2 lung pathology and localization in hACE2-transduced bats

Infection of the lungs was principally localized around the main stem bronchia, with destruction of cilia corresponding with neutrophil infiltration, and lymphoid follicle formation (Figure 4A). Perinuclear SARS-CoV-2 nucleocapsid antigen was readily detected in the lungs of challenged bats for up to 14 days, whereas in unchallenged bats no antigen was detected (Figure 4B). On day 21, all bats were negative for nucleocapsid antigen, suggesting viral clearance. Epithelial, goblet and parenchymal cells had perinuclear staining for nucleocapsid antigen, suggesting these cells were susceptible to the Ad5 transduction of human ACE2 (Figure 4C).

**Figure 4.**
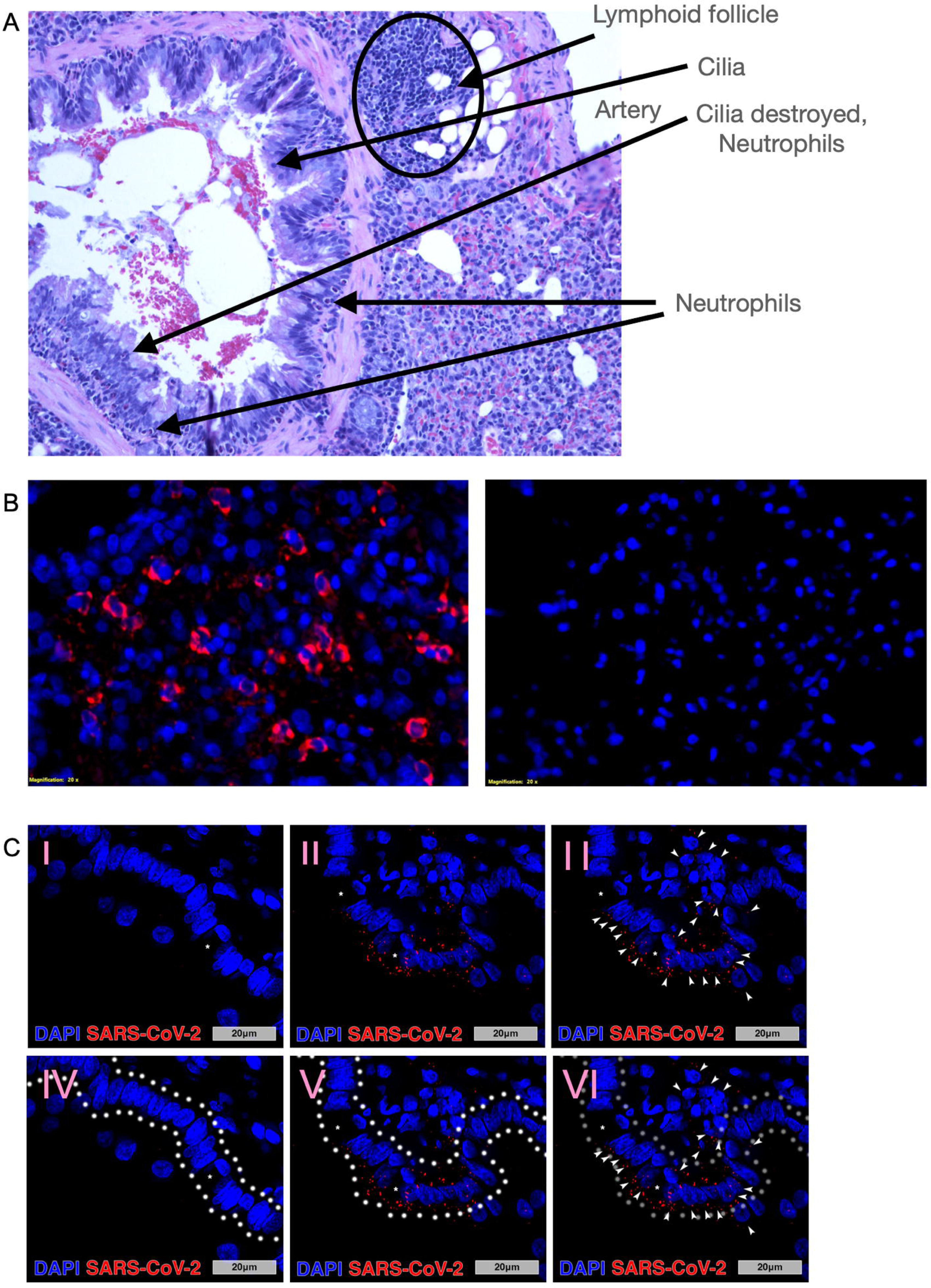
SARS-CoV-2 infection of Jamaican fruit bat lungs transduced with hACE2. **A.** Histopathology of Jamaican fruit bat lungs transduced with Ad5/hACE2 and infected with SARS-CoV-2, 4 DPI. Cilia destruction, neutrophil infiltration, and lymphoid expansion occurred. **B**. Fluorescent imaging of Ad5/hACE2 bat lung sections on day 7 infected with SARS-CoV-2 (left) or uninfected (right) showed perinuclear nucleocapsid antigen staining (red) (200x). **C.** Main stem bronchi (100x) in bats that received Ad5/hACE2 but no SARS-CoV-2 (I, IV) and bats that received Ad5/hACE2 followed by viral infection with SARS-CoV-2 at post-infection day 14 (14 DPI; II, III, V, VI). Asterisks identify goblet cells within the respiratory epithelium. Arrowheads identify cells that contain positive perinuclear SARS-CoV-2 nucleocapsid staining, that are composed of epithelium, goblet cells and cells of the lung parenchyma. White outlines highlight the respiratory epithelial layer (including goblet cells) of the bronchi for reference (IV, V, VI). **D**. SARS-CoV-2 causes loss of cellularity in hACE2-transduced bat lungs. A trend downward was observed on day 2 and was maximal on day 4, with reduced cellularity thereafter that was not significantly different.

### SARS-CoV-2 infection of bats expressing hACE2 leads to diminished cellular population within lung tissue

Total cellular quantification of whole lung tissue in control, Ad5/hACE2, and Ad5/ hACE2/SARS-CoV-2 groups revealed temporal alterations in cellular populations that were dependent upon hACE2 expression and subsequent infection with SARS-CoV-2 (Figure 5). Time course analysis of DAPI^+^ nuclei within the lung parenchyma (Figure 5A) showed reduction in total cellular density over time where the greatest variance from the hACE2 group was observed at 4 DPI when compared to the Ad5/hACE2/SARS-CoV-2 group, and variance from control was detected in the Ad5/hACE2/SARS-CoV-2 group at 10 DPI (Figures 5A, 5B). Driving factors of variance between groupings was attributed to the following: the combination of hACE2, SARS-CoV-2, and time (p=0.0468, F(4,28)=2.768); hACE2 and SARS-CoV-2 (p=0.0002, F(2,28)=11.62) (Figure 5B). Time course assessment of DAPI^+^ bronchial epithelial cells (Figures 5C, 5D) revealed unremarkable changes where trending variance was observed from the combined analysis of hACE2 and SARS-CoV-2 infection (p=0.0514, F(2,28)=3.307).

**Figure 5.**
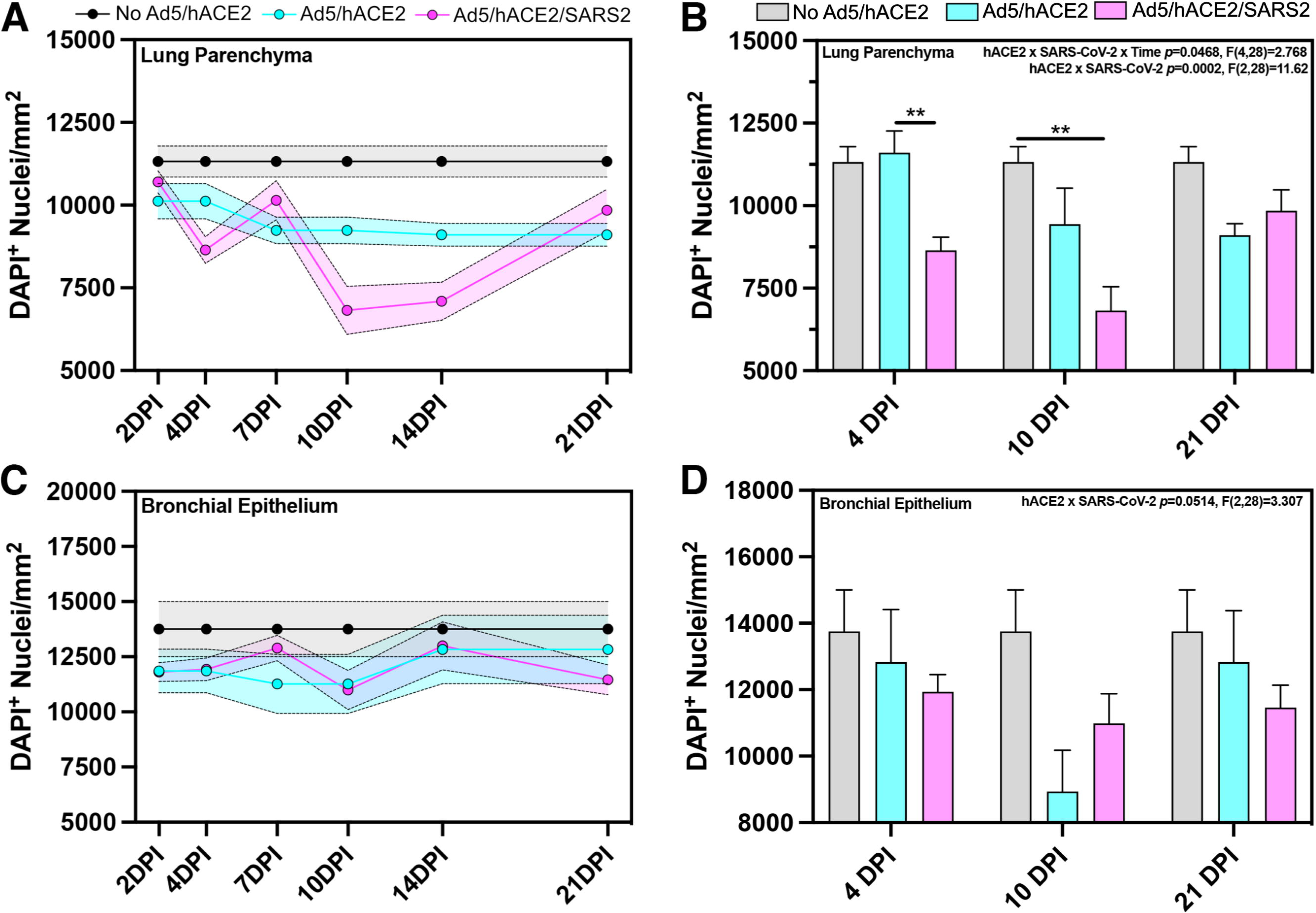
SARS-CoV-2 infection of hACE2 transduced bats leads to reduced numbers of cells in the lung parenchyma that is time dependent. Time course analysis of total DAPI+ cellular populations of whole tissue montage images within the lung parenchyma (A,B) and bronchial epithelium (C,D) of no human ACE2 control bats (grey), hACE2 bats (cyan), and hACE2 + SARS-CoV-2 bats (pink). n=2-5 lung sections per animals/group. **p<0.01

### CD4^+^ T cells responded to SARS-CoV-2 in infected bats

To determine if helper T cells may play a role in bat immune responses during infection, we adopted an activation-induced marker (AIM) test to identify CD4^+^ T cell responses (32–34). This assay relies on increased expression of CD154 on helper T cells that occurs within hours of TCR engagement of MHC-II/peptide and its signal-to-noise ratio is augmented by blocking CD40 on the antigen presenting cells.

We screened several commercially-available anti-CD4 monoclonal antibodies (mAb) for cross reactivity; however, none were useful. Therefore, we generated a mAb (designated 1-D5) using a Jamaican fruit bat CD4 peptide (ENRKVSVVKTRQDRR) that was predicted to be extracellular and solvent-accessible, conjugated to KLH for mouse immunizations (Supplemental Figure 2). Based upon structural homology (Supplemental Figure 3), we determined that commercially available anti-mouse CD40 (Tonbo, clone FGK45) and anti-human CD154 (Tonbo, clone 5C8) monoclonal antibodies were cross reactive with Jamaican fruit bat orthologs (Supplemental Figure 4).

Four bats transduced with Ad5/hACE2 and challenged with SARS-CoV-2 were euthanized 12 days later (Figure 6A). Splenocytes were cultured with or without SARS-CoV-2 nucleocapsid peptide library and anti-CD40 blocking antibody in duplicate for 6 hours. The cells were then co-stained for CD4 and CD154, cells were gated (Supplemental Figure 5), and evaluated for CD154 expression by flow cytometry. Less than 2% of the CD4^+^ cells also expressed CD154, which was significantly elevated on these cells that were stimulated by peptide (p<0.0001) compared to paired samples without peptide stimulation (Figure 6B). Replicate cells from the bats were cultured 24 hours with or without peptide library and RNA extracted for RT-qPCR for T cell gene expression profiling. Three of the four bats had increased expression of several genes; however, one bat did not exhibit changes in gene expression (Figure 6C), suggesting it was not responsive to antigen. Excluding this bat, IL-10, TGFβ, Cxcr4 and CD4 were elevated >2 fold, whereas IL-2, IL-4 and IL-21 expression were substantially downregulated (Figure 6D).

**Figure 6.**
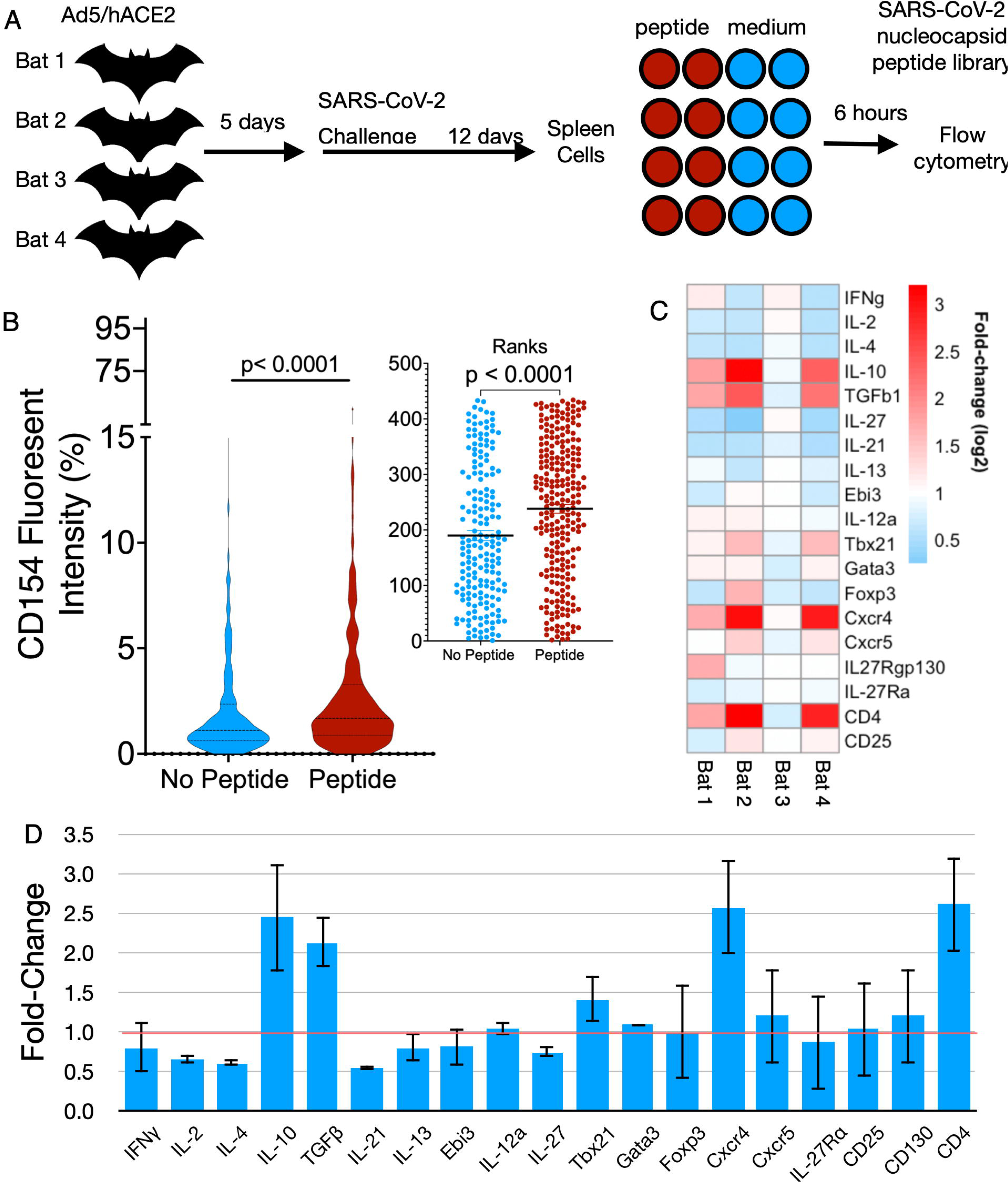
Activation of CD4^+^ Helper T cells in SARS-CoV-2-infected Jamaican fruit bats. (A) Splenocytes from four Ad5/hACE2-transduced bats 12 days post infection with SARS-CoV were cultured with or without SARS-CoV-2 nucleocapsid peptide library for 6 hours in the presence of anti-CD40 blocking antibody. Cells were stained with anti-CD154 and anti-CD4 and analyzed by flow cytometry. (B) In the presence of peptide, CD154 expression was significantly increased on splenocytes (red) compared to no peptide controls (blue) (Wilcoxon rank sum test). (C) Heat map of splenocyte gene expression in response to peptide stimulation determined 3 of the 4 bats expressed a profile consistent with a regulatory T cell response. (D) Gene expression changes in bats 1, 2 and 4.

## Discussion

A significant limitation for studying coronavirus infections of bats is the lack of relevant colonized bat species and the availability of coronaviruses that are naturally hosted by those species. Sarbecoviruses are naturally hosted by insectivorous horseshoe bats (*Rhinolophus* spp.) and there are more than 80 species in the genus. Until such colonies are established, research on bats and coronaviruses will require surrogate bat models, similar to how the laboratory mouse is routinely used as a surrogate model. We established a breeding colony of Jamaican fruit bats in 2006 and have used bats from this colony for a number of infection studies with several viruses, including MERS-CoV (13, 35–37).

The initial challenge study determined that SARS-CoV-2 replicated poorly in Jamaican fruit bats for only a few days without visible signs of disease, is confined to the intestine with detection of virus in the Peyer’s patches and mononuclear cells of the intestinal lamina propria, no apparent shedding in feces, and without a detectable antibody response, even by highly sensitive ELISA to nucleocapsid antigen. The lack of detectable virus in the lungs is consistent with what is known about coronavirus circulation in wild bat populations, where virus is typically found in the intestines and shed through feces. One possible explanation for this is that very low levels of endogenous ACE2 mRNA was detected in Jamaican fruit bat lungs. The only other experimental SARS-CoV-2 challenges of bats have yielded varying results. An experimental challenge of Egyptian fruit bats led to a limited infection without visible signs of disease, but with detectable virus in several tissues, including lungs and lymph nodes, and transmission to one contact bat (28). Challenge of big brown bats did not lead to infection (29); however, two studies of Mexican/Brazilian free-tail bats were contradictory, with one suggesting they were susceptible but not the other (30, 31). ACE2 residues that are thought to be important in SARS-CoV-2 spike binding (38, 39) are more conserved in Egyptian fruit bats (16/20 residues with human), substantially less in big brown bats and Mexican/Brazilian free-tailed bats (11/20 residues), with Jamaican fruit bats intermediate (13/20 residues) (Supplemental Table 1). The conservation of these ACE2 residues varies substantially among several horseshoe bat species; thus, it may be that certain horseshoe bat species may favor evolution of sarbecoviruses that are more likely to infect humans based upon similarity of ACE2 contact residues. If so, then such conservation may be predictive of which horseshoe bat species should be monitored for sarbecoviruses. Currently, only ten horseshoe bat species ACE2 sequences are available in GenBank, which is a fraction of the number species in the genus. Field studies to determine sequences of ACE2 from other horseshoe bats may be useful for identifying potential sarbecoviruses with pandemic potential.

It may be that SARS-CoV-2 spike protein, which mediates cellular attachment to ACE2, is substantially different compared to its ancestral virus because of the likely route of passage through one or more intermediate bridge hosts before spillover into humans, and subsequent adaptations as it transmitted among humans. As SARS-CoV-2 has continued to circulate among humans, new variants that have emerged are likely to be more divergent from the ancestral virus, such that SARS-CoV-2 may now be more of a “human virus” than a “bat virus.” Ideally, the original bat virus that spilled over into the bridge host(s) is required to fully understand the biology of the virus. However, it is unlikely the virus will ever be found because by now it may have gone extinct or has undergone substantial recombination with other sarbecoviruses, a common occurrence of coronaviruses (1, 40), such that it no longer exists in nature. The earliest SARS-CoV-2 isolate available to us was WA1, which was isolated from a patient in Washington state, USA. It may be that earlier isolates, such as the initial Wuhan isolates, may have behaved differently in Jamaican fruit bats. Regardless, the results presented here suggest that SARS-CoV-2 has limited ability to bind Jamaican fruit bat ACE2 (AjACE2) (poor susceptibility), which is consistent with a report demonstrating spike binding to AjACE2 (41), it is unable to replicate in Jamaican fruit bat cells (poor permissibility), the innate response controls infection, or a combination of these features.

Several adenovirus serotypes use the coxsackievirus and adenovirus receptor (CAR, *Cxadr*) for cellular entry. We determined that Jamaican fruit bat CAR protein shares 86% identity and 92% similarity with the mouse ortholog, suggesting it could be used for adenovirus transduction of human ACE2. To address this possibility, we adopted an approach previously developed for the assessment of SARS-CoV-1 and SARS-CoV-2 infection of laboratory mice, in which a replication-defective serotype 5 adenovirus (Ad5), which uses CAR for cellular entry, was generated that expresses human ACE2 (Ad5/hACE2) (32–34, 42). Upon transduction of Jamaican fruit bat primary kidney epithelial (Ajk) cells with Ad5/hACE2/eGFP, sustained fluorescence was observed for several days. However, inoculation of the cells with SARS-CoV-2 did not lead to detectable virus replication, suggesting the cells are not permissive. It is also possible that the cells lacked a suitable protease, such as TMPRSS2, that can facilitate SARS-CoV-2 penetration of the plasma membrane, or that adenovirus transduction activated antiviral pathways in the cells that restricted SARS-CoV-2 replication. Alternatively, SARS-CoV-2 can infect certain cells, such as Vero E6, via endosomal entry where cathepsin L facilitates endosomal escape of the genomic RNA into the cytoplasm (43). A recent study determined that cell lines from several insectivorous bat species were not permissive to SARS-CoV-2 replication despite expression of endogenous bat ACE2 on some of these cells. One of the cell lines tested was from a patagium biopsy of a greater horseshoe bat (*Rhinolophus ferrumequinum*), a member of the genus that harbors sarbecoviruses (1); however, it was unclear whether these cells expressed endogenous ACE2, or whether a lack of sufficient RfACE2 binding residues may impair spike binding (12/20 residues, Supplemental Table 1), and attempts to transduce this cell line with human ACE2 failed (44). Transduction of human ACE2 led to low-level, nonproductive SARS-CoV-2 infection in four of the other bat cell lines, but none were of kidney origin like Ajk cells. One of the lines appeared to produce virus particles; however, electron microscopy indicated the viruses were unable to complete exocytosis. The authors postulated that tetherin, which appears to be under strong selective pressure in bats (45), may account for this observation. In the other cells, expression of *Oas1* and *Ifih1* (MDA-5) transcripts were elevated during SARS-CoV-2 infection and may have impaired virus replication. Thus, our results are consistent with these findings and may explain the lack of viral shedding in Ad5/ hACE2 bat lungs (Figure 3B).

Despite the discouraging results of the Ajk cells, we proceeded with hACE2 transduction of cells in the lungs of Jamaican fruit bats to determine susceptibility. Transduced bats challenged with 10^5^ TCID_50_ equivalents of SARS-CoV-2 WA1 did not show visible signs of disease and they did not lose weight (Figure 3D). BALB/c mice similarly transduced with hACE2 and challenged with SARS-CoV-2 exhibited significant weight loss of 10-25% during the first week but recovered thereafter (32). It possible that the detected virus (Figure 3B) represents residual input virus; Hassan et al. detected low levels of viral RNA in the lungs of SARS-CoV-2 challenged BALB/c mice that had not been transduced with hACE2 (32). Regardless, clear pathology was present in hACE-transduced and infected bat lungs (Figure 4A), antigen was readily detected in the main stem bronchi of challenged bats for up to 14 days, and clear but low-titer antibody responses occurred beginning on day 7 (Figure 3C, Supplemental Figure 1). This is consistent with work by us and others showing that bats typically produce low-titer antibody responses to viral infections (28, 35, 37, 46–50). We cannot exclude the possibility that adenovirus transduction elicited an innate response that could have augmented control of SARS-CoV-2 infection; however, mouse studies have shown innate stimulation from transduction subsides within a day of inoculation with the vector (51).

Total cellular quantification conducted in the bronchial epithelium and parenchyma revealed significant decreases in the amount of total cellular populations at 4 DPI and 10 DPI that recovered to baseline at 21 DPI in animals that were transfected with hACE2 adenovirus and subsequently challenged with SARS-CoV-2 (Figure 5). Notably, SARS-CoV-2 infection in other animal models and reports of clinical cases showed compelling histopathological evidence that SARS-CoV-2 infection leads to immune cell infiltration of the lung parenchyma and collapse of bronchial airways, evident through loss of bronchial epithelium (52–55). The recruitment of immune cells to the lung in SARS-CoV-2 infection can occur as early as 2 DPI and is often progressive and results in permeant lung damage, severe disease, fibrosis, and/or death (54). However, the reduction in total cellular populations within the lung parenchyma in concert with the retention of bronchial epithelium in bats transduced with hACE2 and then challenged with SARS-CoV-2 shows that these bats become susceptible to viral infection within the lungs only if hACE2 is present. Despite this, the apparent pathology does not appear to be as severe as in other mammalian model systems or clinical patients (52–55). These results provide compelling evidence that although expression of hACE2 in bats allowed infection, the immune response of the bats likely mitigated severe pathological outcome and hypercellularity within the lungs.

We also determined that CD4^+^ helper T cells responded to SARS-CoV-2 infection in the hACE2 bats. Antigen-reactive T cells typically are rare in infected animals, usually fewer than 2% of T cells even days or weeks after infection, making assessment of ex vivo T cell responses challenging. However, because CD154 is an activation marker of helper T cells, its expression can be used to identify the rare helper T cells reactive to antigen by flow cytometry (56, 57). We found a significant increase in CD154 (CD40L) expression on CD4^+^ helper T cells when stimulated with a SARS-CoV-2 nucleocapsid peptide library (Figure 6B). In humans, abundant CD4^+^CD154^+^ T cells are associated with severe COVID-19, particularly in T cells producing abundant proinflammatory cytokines, including interferon-γ (IFNγ), tumor necrosis factor, interleukin-2 (IL-2) and/or interleukin-21 (IL-21) (58, 59). In the splenocytes of 3 of 4 bats cultured 24 hours with nucleocapsid peptide library antigen, IFNγ, IL-2 nor IL-21 transcripts were elevated, whereas IL-10 and TGFβ were elevated more than two-fold (Figures 5C/D), despite the low frequency (<2%) of antigen-specific helper T cells in the splenocyte cultures. These two cytokines are typically produced by regulatory T cell subsets that suppress inflammatory responses (60). Expression of Th1 transcripts IL-2 and IFNγ, Th2 transcripts IL-4 and IL-13, and T follicular helper (Tfh) IL-21 transcripts were lower than basal level expression. Fox-p3 and Cxcr5 expression were not substantially elevated, which could be explained by low levels of expression of this transcription factor and cell surface receptor, respectively, in the few antigen-reactive T cells (i.e., low signal-to-noise ratios). Identification or development of monoclonal antibodies for these bat orthologs could enhance their detection using flow cytometery, particularly to determine whether the Treg-like cells are expressing both IL-10 and TGFβ, or if there are two populations expressing each cytokine. Transcripts of helper T cell markers CD4 and Cxcr4, which is substantially elevated in patients with severe COVID-19 (61), were both elevated, suggesting that Cxcr4 activity may be less in bats compared to humans, or it may have a different role in bats. This study also determined that Treg cell responses during COVID-19 may lead to more severe outcomes in humans, which is seemingly contradictory considering IFNγ- and IL-21-expressing T cells appeared to contribute to immunopathology (59). Thus, there appear to be similar responses, yet clear differences in helper T cell behaviors in bats and humans infected with SARS-CoV-2. The T cells from bat 3 did not have elevated expression of any gene in the qPCR array (Figure 5C). Repeated testing of the duplicate samples from this bat yielded similar results; thus, we are confident the RNA was properly extracted. It is also possible that the nucleocapsid peptide library was not added when the splenocytes were plated; however, because each T cell culture from each bat was performed in duplicate we think this is unlikely because replicates from both had similar Ct values. It may be that other genes not on the qPCR array were expressed by these T cells; RNA-Seq could clarify this question.

Helper T cells are instrumental for many effector functions and provide the highest level of command and control of immune responses. Indeed, engagement of Th cell CD154 by B cell CD40 leads to T cell help, including IL-21 production from Tfh cells in germinal centers (62), for class switching and somatic hypermutation/affinity maturation. In the present study, we only detected low titer antibody responses in the bats, even at 21 days post infection, and the lack of IL-21 expression may be one reason why antibody titers, which are principally a function of affinity maturation, are poor. In our previous experimental infection studies of Tacaribe virus-infected Jamaican fruit bats, which causes a fatal disease, RNA-Seq analysis identified genes required for somatic hypermutation in spleens, including activation-induced cytidine deaminase; however, they were not differentially expressed in infected bats (36). Thus, it may be that bats have such robust innate immune responses that they are less dependent on antibody responses. However, prior to the present work, no studies have been published describing virus-specific helper or cytotoxic T cells in bats, principally because of a lack of tools for the study of bat adaptive immunity; thus, it is unclear what roles bat T cells play in immunity to viruses. It may be that bats have a greater dependency on regulatory T cells that could temper inflammatory pathology, which is consistent with the observation that bats typically have low or no inflammatory responses during viral infections (35, 46, 63, 64). This also parallels what occurs in rodent reservoirs of hantaviruses, in which TGFβ-expressing regulatory T cells predominate to what are otherwise innocuous infections (65, 66). Further studies requiring the development of new reagents and methods for bats will be needed to address these questions, particularly studies for examining individual T cells (e.g., T cell culturing, flow cytometry, single-cell RNA-Seq, etc.).

There are limitations of this study that should be addressed in future research. We are currently unable to assess cytotoxic T cell responses in bats. AIM tests have been developed for cytotoxic T lymphocytes by assessing elevated CD134 expression on CD8^+^ T cells (67). However, we have been unable to identify a monoclonal antibody that recognizes Jamaican fruit bat CD8, thus one will need to be developed. Identification of cross-reactive antibodies to CD134 will also be necessary; otherwise, monoclonal antibodies to these will need to be generated. It will also be necessary to examine the kinetics of the T cell response using various time points and larger sample sizes for AIM testing and cytokine gene expression. Bats are of low fecundity and female Jamaican fruit bats give birth to one pup at 6-month intervals, limiting their availability for such studies.

The findings here also show that serotype 5 adenovirus vectors are important tools for addressing viral infections of bats, including SARS-CoV-2 variants. It is likely that SARS-CoV-2 originated in horseshoe bats, thus it should be possible to address questions of SARS-CoV-2 and other sarbecovirus susceptibility and infection of Jamaican fruit bats transduced with ACE2 from horseshoe bat species. Although work with cells expressing various bat ACE2 proteins have been performed (41), a better understanding of how these viruses behave in bats, particularly adaptive immune responses which require live bats, could shed light on the biology of coronaviruses. A limitation of adenovirus use is that gene transduction is confined to the respiratory tract and not the intestines, where natural bat coronaviruses are typically found. Nonetheless, this approach offers a means to more fully dissect the role of a variety of viral infections of bats and how the adaptive immune system responds to those viruses.

## Materials and Methods

### Jamaican fruit bats

This study was performed with approval from the Colorado State University Institutional Animal Care and Use Committee (protocol 1787). Bats of both sexes were used; however, more males than females were used because the low fecundity (one pup at 6 month intervals for reproductive females). The bat colony is maintained in a free-flight facility; however, Infected bats were confined to bird cages under BSL-3 containment upon challenge with SARS-CoV-2.

### Generation of anti-Jamaican fruit bat CD4 monoclonal antibody

The Jamaican fruit bat CD4 protein was submitted to Phyre2 molecular modeling server and the generated structure file provided several candidate solvent-accessible peptides. An extracellular membrane-proximal sequence from residues 236-277 (ENRKVSVVKTRQDRR) with an N-terminal cysteine was generated and covalently linked to KLH (Genscript). BALB/c mice were intraperitoneally-immunized with 25 µg of the KLH-peptide conjugate emulsified in incomplete Freund’s adjuvant (IFA) and boosted at 1 month intervals (in IFA) for 3 months. Sera from mice were reactive to Jamaican fruit bat splenocytes. A final boost in PBS was administered and 4 days later splenocytes were fused with Sp2/10 Ag14 cells (ATCC CRL-1581) and cultured overnight prior to concurrent cloning and HAT selection on methylcellulose plates (ClonaCell-HY, StemCell Technologies). After 14 days of incubation at 37°C, clones were picked for expansion. Supernatants were screened by flow cytometry to identify reactive monoclonal antibodies. Several were identified and one, 1-D5 was selected for fluorescent labeling.

### Identification of cross reactive anti-CD40 and anti-CD154 antibodies: in silico protein homology

Jamaican fruit bat CD40 (NCBI, XP_037020498) and CD154 (NCBI, XP_037012946) polypeptide sequences were submitted to Phyre2 molecular modeling server to generate PDB files (68). The generated Jamaican fruit bat protein PDB structures, mouse CD40 (UniProt, P27512), and human CD154 (UniProt, P29965) PDB files were imported into Geneious Prime for the assessment of protein homology by Blocks Substitution Matrix 62 (BLOSUM62) alignment (69). The three-dimensional structures were then annotated for the transmembrane domains and identical amino acid residues to identify putative conserved regions to which commercially-available monoclonal antibodies might bind to Jamaican fruit bat orthologs.

### Identification of cross reactive anti-CD40 and anti-CD154 antibodies: flow cytometry

One naïve male Jamaican fruit bat was euthanized and a single-cell suspension of its spleen was generated by mechanical disassociation in cold 10 ml PBS with 5 mM EDTA by passage through a 70µm cell strainer. Splenocytes were then placed in a 15 ml conical tube and centrifuged at 4°C for 5 minutes at 350 g and supernatant was decanted. Splenocytes were then resuspended in cold 10 ml ammonium chloride solution (150mM NH_4_Cl, 10mM NaHCO_3_, 10mM EDTA in cell culture grade water) and incubated for 10 minutes on an orbital rotator to lyse red blood cells. The cells were washed and resuspended in 10 ml PBS, 5 mM EDTA splenocytes were then counted using a hemocytometer and trypan blue and resuspended in cryopreservation media (90% FBS, 10% DMSO) at 5×10^6^ cells/ml, and 1 ml aliquots dispensed into cryovials. Cryovials were then stored in −80°C for 2 days, then transferred to liquid nitrogen.

One cryovial was removed from liquid nitrogen, thawed in a 37°C water bath, transferred to 15ml conical tube and washed with 10ml PBS, 5 mM EDTA. 10^6^ cells were transferred to wells of a 96 well v-bottom plate for the unstained control. Splenocytes were washed and resuspended in 5 ml of a 1:5,000 dilution of primary amine viability dye (Ghost DyeTM Red 780) in PBS with 5 mM EDTA solution and incubated in the dark at 4°C for 30 minutes. Splenocytes were then centrifuged at 4°C for 5 minutes at 350 g and supernatant was decanted. Splenocytes were then washed in 10ml FACS buffer (PBS, 1% BSA, 0.5% sodium azide, 5 mM EDTA) centrifuged at 4°C for 5 minutes at 350 g and supernatant was decanted. Cells were then resuspended in FACS buffer, and 1 million splenocytes were added 96 well v-bottom plate per test sample. Splenocytes were then centrifuged at 4°C for 3 minutes at 350 g and supernatant was decanted. Splenocytes were then resuspended in 100µl Fc receptor blocker (Innovex, ready-to-use) and incubated for 30 minutes at 4°C in the dark. Splenocytes were then centrifuged at 4°C for 3 minutes at 350g and supernatant was decanted. Test samples were then resuspended in a 1:20 dilution of anti-mouse CD40 PE (Tonbo, clone FGK45), or anti-human CD154 PE (Tonbo, clone 5C8) in FACS buffer and incubated for 30 minutes at 4°C in the dark. Splenocytes were then centrifuged at 4°C for 5 minutes at 350 g and supernatant was decanted. Splenocytes were washed 2x with 150µl FACS buffer centrifuged at 4°C for 5 minutes at 350g and supernatant was decanted each time. Splenocytes were then resuspended in 100µl FACS buffer. A 4 laser Cytek Auora (16V-14B-10YG-8R) cytometer was used to acquire the FCS files. The unmixed FCS files were then analyzed using FlowJo 10.8.1.

### SARS-CoV-2 infections

The WA1 isolate of SARS-CoV-2 was used for this study and was obtained from BEI Resources and passaged twice in Vero E6 cells to generate a stock. Intranasal inoculations were performed while holding the bats in a contralateral position to ensure delivery to the trachea and the esophagus (i.e., lungs and GI tracts) with 10^5^ TCID_50_ equivalents per bat in 50 µl in sterile PBS. Oral and rectal swabs were collected from bats on days 2, 4, 7, 10, 14 and 21 post-inoculation (PI) for detection of virus by qPCR. Groups of bats were euthanized on days 2, 4, 7, 10, 14 and 21 PI for necropsy with portions of lungs and intestine collected and frozen for viral RNA and virus isolation, and the remaining tissues collected in buffered formalin.

### Virus detection

For virus isolation, frozen lungs and intestines were homogenized with a Tissue Lyser II in 10% FBS DMEM, centrifuged at 8,000g for 5 min, then passed through a 0.2 µm filter to remove bacterial contaminants. Log_10_ dilution series were made for each sample and added to replicate 96 wells of Vero E6 cells and scored for CPE 5 days later to calculate TCID_50_. For qPCR, RNA was extracted from frozen tissues (Qiagen RNEasy kit) and TCID_50_ equivalents determined using a SARS-CoV-2 E gene detection kit (Promega) based upon the Berlin primer/ probe set (70).

### Adenovirus transduction of human ACE2

A serotype 5 defective adenovirus encoding the human ACE2 gene (Ad5CMVhACE2) purchased from the University of Iowa Viral Vector Core was used for this study. The E1 gene is replaced with human ACE2, rendering the virus incapable of replication. For in vitro studies, Jamaican fruit bat Ajk cells were inoculated with 1 MOI of a GFP-expressing virus (Ad5CMVACE2-IRESeGFP) for 1 hour, followed by removal of inoculum and 2x washing in 2% FBS-DMEM. For SARS-CoV-2 challenge, cells were inoculated with virus without GFP (Ad5CMVhACE2) as above and at 48 hours inoculated with 0.1 MOI of SARS-CoV-2. Supernatants were collected at 1 hr and daily thereafter for titration on Vero E6 cells. For in vivo experiments, bats were intranasally inoculated with 2.5×10^8^ pfu in 50 µl of serum-free DMEM. SARS-CoV-2 challenge was performed 5 days later.

### Histopathology and Immunohistochemistry

Tissues and carcasses with open abdomens were collected in 10% neutral buffered formalin for 3 days prior to removal from the BSL-3 and transferred to the Colorado State University Diagnostic Laboratories for trimming. Oral cavity, salivary glands, olfactory bulb, cerebrum, cerebellum, and brain stem were thoroughly inspected for gross lesions. Decalcified skulls and visceral organs were processed, embedded in paraffin wax and 4–5 µm sections were stained with hematoxylin and eosin for blinded evaluation by the pathologist using Nikon i80 microscope (Nikon Microscopy).

Sections were stained using ultraView universal alkaline phosphatase red detection kit. Heat-induced epitope retrieval was performed on a Leica Bond-III IHC automated stainer using Bond Epitope Retrieval solution for 20 minutes. Viral nucleocapsid antigen was detected with a purified rabbit polyclonal antibody. Labeling was performed on an automated staining platform. Fast Red was used as chromogen and slides were counterstained with hematoxylin. Immunoreactions were visualized by a single pathologist in a blinded fashion. In all cases, normal and reactive mouse brain sections incubated with primary antibodies was used as a positive immunohistochemical control. Negative controls were incubated in diluent consisting of Tris-buffered saline with carrier protein and homologous nonimmune sera. All sequential steps of the immunostaining procedure were performed on negative controls following incubation.

### Fluorescent microscopy

Paraffin embedded tissue sections were stained for SARS-CoV-2 nucleocapsid protein (1:500) using a Leica Bond RXm automated staining instrument following permeabilization using 0.01% Triton X diluted in Tris-buffered saline (TBS). Blocking was performed with 1% donkey serum diluted in TBS. Sections were stained for DAPI (Sigma) and mounted on glass coverslips in ProLong Gold Antifade mounting medium and stored at ambient temperature until imaging. Images were captured using an Olympus BX63 fluorescence microscope equipped with a motorized stage and Hamamatsu ORCA-flash 4.0 LT CCD camera. Images were collected and regions of interest quantified with Olympus cellSens software (v 1.18) using an Olympus X-line apochromat 10X (0.40 N.A.), 20X (0.8 N.A.) or 40X (0.95 N.A.) air objectives, or Uplan Flour X100 oil immersion (1.3 N.A.) objective.

### Automated high-throughput immunofluorescence staining and imaging of tissue sections

Paraffin embedded lung tissue was sectioned at 10 µm thickness and mounted onto poly-ionic slides (Colorado State University Diagnostic Laboratory). Tissue sections were deparaffinized and immunofluorescently labeled using a Leica Bond RXm automated robotic staining system. Antigen retrieval was performed by using Bond Epitope Retrieval Solution 2 for 20 minutes in conjunction with heat application. Sections were then incubated with primary antibodies diluted in 0.1% triton-X containing tris-buffered saline (TBS): rabbit anti-SARS-CoV-2 nucleocapsid protein (Novus Biologicals; 1:100) and goat anti-human angiotensin converting enzyme 2 (ACE2; R&D Systems; 1:500). Appropriate species-specific secondary antibodies conjugated to AlexaFluor555 and/or AlexaFluor647 were applied for fluorescent detection. Sections were stained with DAPI (Sigma, 1:5,000) and mounted on glass coverslips in ProLong Gold Antifade hard set mounting medium and stored at 4°C until time of imaging.

### Artificial intelligence neural network detection and quantification of total nuclei in whole tissue sections

Whole lung section montage images were acquired on a fully automated motorized stage scanning VS200 microscope (Evident, Waltham, MA USA) equipped with a Hamamatsu ORCA-Fusion camera (Hamamatsu Photonics, Shizuoka, Japan) and imaged with at 200X resolution using an extended apochromat X-line 20X air (0.8 N.A.) using Olympus CellSens (v.3.2). All slides were stained and imaged simultaneously using consistent exposure times and CCD gain and binning parameters to reduce variability in intensity measurements. Images were then imported into Visiopharm Software (v.2022.09.0.12417) and the immunofluorescence nuclear detection application was modified to accommodate intensity range and average area of cells within the lung sections. Regions of interest (ROIs) were generated around whole lung sections based on DAPI positive staining intensity, where total nuclei detected within the tissue was normalized to the total area of whole tissue to minimize variation observed between differing sizes of sectioned lung.

### Serum IgG ELISA

The SARS-CoV-2 spike (S) ectodomain containing the prefusion stabilizing hexapro mutations (71) and a mutated furin cleavage site was produced and purified from Expi293 cells as previously described (72). The SARS-CoV-2 S protein was diluted to 0.003 mg/ mL in PBS and used to coat 384-well Nunc Maxisorp plates (Thermo Fisher) overnight at room temperature. The plates were slapped dry and blocked for 1 hour at 37°C using Casein in PBS (Thermo Fisher). Following blocking, the plates were again slapped dry and sera from infected bats was added to the plates beginning at a 1:30 dilution in TBST followed by a 1:3 serial dilution thereafter. Plates were incubated for 1 hour at 37°C, slapped dry, and washed four times with TBST. Recombinant protein A/G conjugated to HRP (Thermo Fisher) diluted 1:500 in TBST was added each well and the plates were incubated for 1 hour at 37°C then washed four times with TBST. To measure binding titers, TMB Microwell Peroxidase (Seracare) was added to each well and, after 2 minutes, the reaction was quenched with 1 N HCl. The absorbance at 450 nm was measured using a BioTek Neo2 plate reader and analyzed in GraphPad Prism 9 with ED_50_ values being determined using a four-parameter logistic regression model. Two biological replicates were performed for each sample.

### Neutralization assays

VSV particles pseudotyped with the SARS-CoV-2 S protein harboring the D614G mutation were produced as previously described (73, 74). To perform the neutralization assays, 20,000 Vero-TMPRSS2 cells were seeded into each well of a 96-well plate and grown overnight until they reached approximately 80-90% confluency. Sera from infected bats were diluted in DMEM (Gibco) beginning at a 1:10 dilution followed by a 1:3 serial dilution thereafter. The diluted sera were then incubated for 30 minutes at room temperature with SARS-CoV-2 S pseudotyped VSV diluted 1:25 in DMEM along with an anti-VSV-G antibody (I1-mouse hybridoma supernatant diluted 1:25, from CRL-2700, ATCC) to block entry of any residual VSV G pseudotyped virus. The Vero-TMPRSS2 cells were washed three times with DMEM and the virus-sera mixture was added to the cells. Two hours after infection, an equal volume of DMEM supplemented with 20% FBS and 2% PenStrep was added to each well. The cells were incubated for 20-24 hours after which ONE-GloEX (Promega) was added to each well and the plates were incubated for 5 minutes. Luminescence values measured in relative light units (RLU) were recorded using a BioTek Neo2 plate reader. The values were normalized in GraphPad Prism 9 with the RLU values from uninfected cells to determine 0% infectivity and the RLU values from cells infected with pseudovirus only to determine 100% infectivity. ID50 values were determined from the normalized data using a [inhibitor] vs. normalized response – variable slope model. Neutralization assays were performed in duplicate and replicated with two distinct batches of pseudovirus.

### Activation-induced marker test

Five bats were transduced with hACE2 and 5 days later inoculated with SARS-CoV-2. Twelve days later, the bats were euthanized and spleens collected to make single cell suspensions. Red blood cells were lysed with ammonium chloride buffer and 10^6^ splenocytes from each bat were cultured with 1 µg of SARS-CoV-2 PepTivator® SARS-CoV-2 Prot_N peptide library (15mers with 12 residue overlaps, Miltenyi) for 6 hours in 5% FBS Clicks medium (Fujifilm/Irvine Scientific) and 1 µg of anti-mouse CD40 monoclonal antibody (FGK45, Tonbo/Cytek, to block CD154 ligation) at 37° C in duplicate. Identical wells were set up in duplicate without peptide library to provide baseline marker levels. Cells were collected, washed and stained with anti-Jamaican fruit bat CD4 and anti-human CD154 (5C8, Tonbo/Cytek). Cells were fixed in paraformaldehyde and examined by flow cytometry. Gating on CD4 was performed and the mean fluorescent intensity (MFI) of CD154 was determined. MFI were evaluated between groups using Wilcoxon signed-rank test.

### Immune gene expression profiling

Identical cultures described in the AIM test were examined for immune gene expression. Splenocytes (5×10^6^) were cultured in 24 well plates with 5 µg of SARS-CoV-2 PepTivator® SARS-CoV-2 Prot_N peptide library for 24 hours. Identical cultures without peptide library were used to determine baseline gene expression. Duplicate splenocyte cultures were examined for changes in gene expression by SYBR Green qPCR array as previously described (21, 66, 75). Briefly, total RNA was extracted from the cells (RNEasy kit, Qiagen) and cDNA produced (QuantiTech RT, Qiagen). Primers for Jamaican fruit bat genes (Supplemental Table 2) and qPCR was performed (QuantiTech SYBR Green, Qiagen). Within sample gene normalization was performed on Rps18 (ΔCq), and fold-chage determined by comparing each gene from the peptide-stimulated cells to the unstimulated cells from the same bat (ΔΔCq).

## Acknowledgments

Funding for this study was provided through grants from the National Institute of Allergy and Infectious Diseases (NIAID) R01 AI140442 (TS, SRW), the National Science Foundation (2033260, TS; 2020297257, BB), National Bio and Agro-Defense Facility (NBAF) Transition Fund from the State of Kansas (JAR), the MCB Core of the Center on Emerging and Zoonotic Infectious Diseases (CEZID) of the National Institutes of General Medical Sciences under award number P20GM130448 (JAR), the NIAID Centers of Excellence for Influenza Research and Surveillance under contract number HHSN 272201400006C (JAR) and the NIAID supported Center of Excellence for Influenza Research and Response (CEIRR) under contract number 75N93021C00016 (JAR), NIAID DP1AI158186 and 75N93022C00036 to DV, the National Institute of Health Cellular and Molecular Biology Training Grant T32GM007270 to AA, a Pew Biomedical Scholars Award (DV), an Investigators in the Pathogenesis of Infectious Disease Awards from the Burroughs Wellcome Fund (DV), Fast Grants (DV). DV is an investigator of the Howard Hughes Medical Institute.

