## Supplemental Figure 2 for "Regulatory T Cell-like Response to SARS-CoV-2 in Jamaican Fruit Bats (*Artibeus jamaicensis*) Transduced with Human ACE2"

**Supplemental Figure 2.** (A) Predicted extracellular structure of Jamaican fruit bat CD4 with immunizing peptide in orange, and (B) flow cytometric staining of naive bat splenocytes with monoclonal antibody 1-D5.

A B

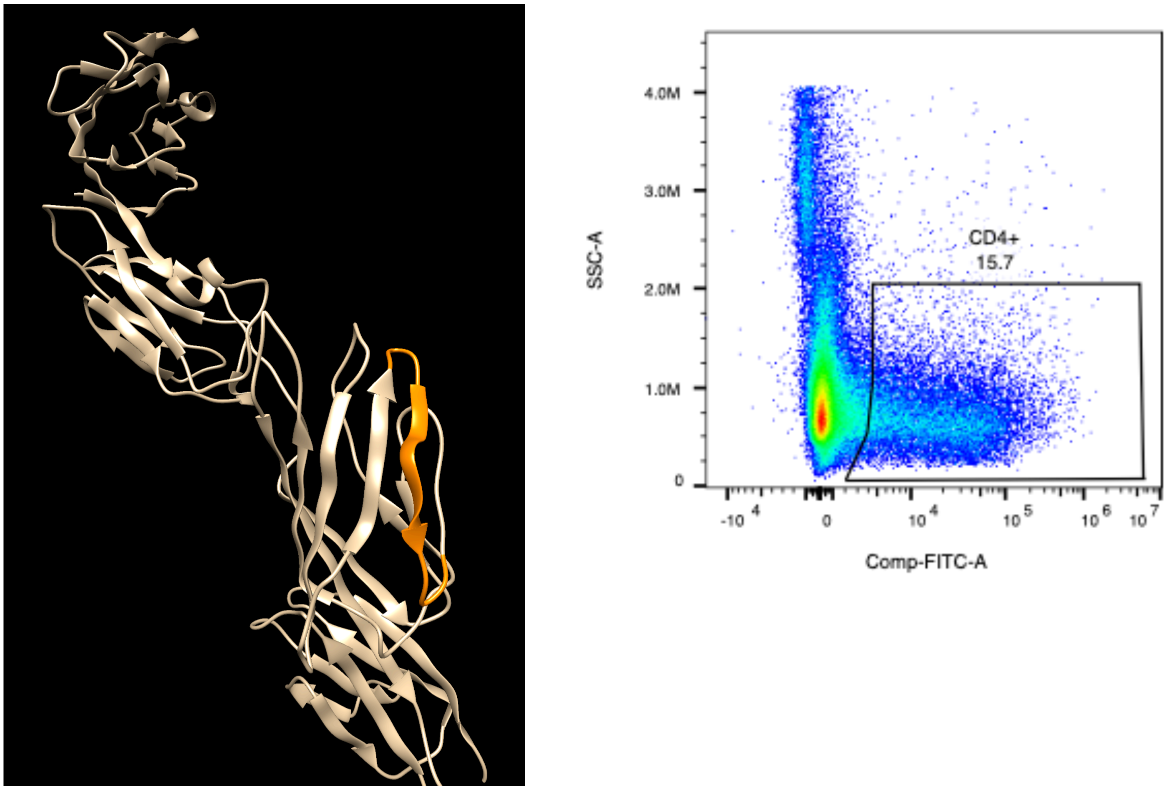
