## Supplemental Figure 3 for "Regulatory T Cell-like Response to SARS-CoV-2 in Jamaican Fruit Bats (*Artibeus jamaicensis*) Transduced with Human ACE2"

**Supplemental Figure 3**. Protein homology of Jamaican fruit bat, mouse, and human CD40 and CD154. CD40 protein structures mouse (top left) and Jamaican fruit bat (top right). CD154 protein structures human (bottom left) and Jamaican fruit bat (bottom right). Protein structures are represented with signal peptides removed and transmembrane domains (magenta) highlighted for orientation. Jamaican fruit bat CD40 and mouse CD40 protein alignment identified 90 identical extracellular sites (red) or 53.39% identical extracellular domains with a BLOSUM62 value of 71.3%. Jamaican fruit bat CD40 and mouse CD40 protein alignment identified 39 identical cytoplasmic sites (cyan) or 72% identical cytoplasmic domains with a BLOSUM62 value of 84%. Jamaican fruit bat CD154 and human CD154 protein alignment identified 190 identical extracellular sites (red) or 88.4% identical extracellular domains with a BLOSUM62 value of 92.1%. Jamaican fruit bat CD154 and mouse CD154 protein alignment identified 18 identical cytoplasmic sites (cyan) or 61.9% identical cytoplasmic domains with a BLOSUM62 value of 73%.

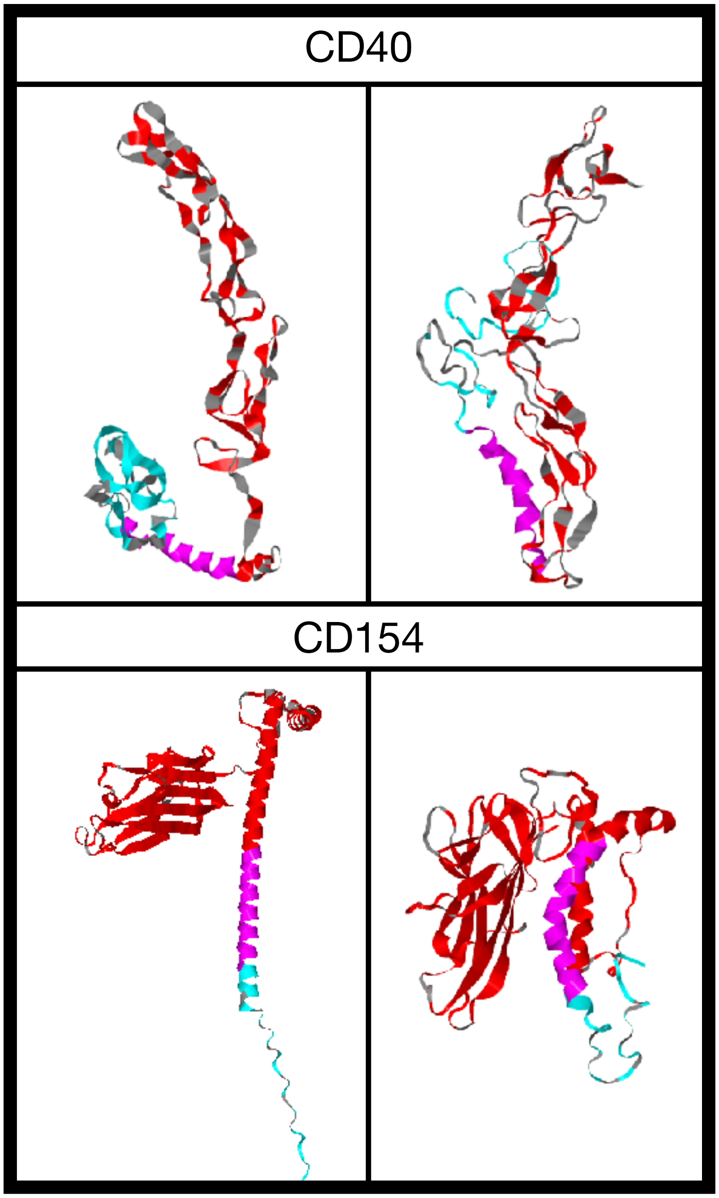
