## Supplemental Figure 4 for "Regulatory T Cell-like Response to SARS-CoV-2 in Jamaican Fruit Bats (*Artibeus jamaicensis*) Transduced with Human ACE2"

**Supplemental Figure 4**. Anti-mouse CD40 (FGK45) and anti-human CD154 (5C8) PE conjugated antibody staining of Jamaican fruit bat splenocytes. Gating strategy: Intact > Live > Single Cells > AF- > PE+. Anti-mouse CD40 cross reactive antibody demonstrated 20.7% positive splenocytes. Anti-human CD154 cross reactive antibody demonstrated 1.69% positive splenocytes.

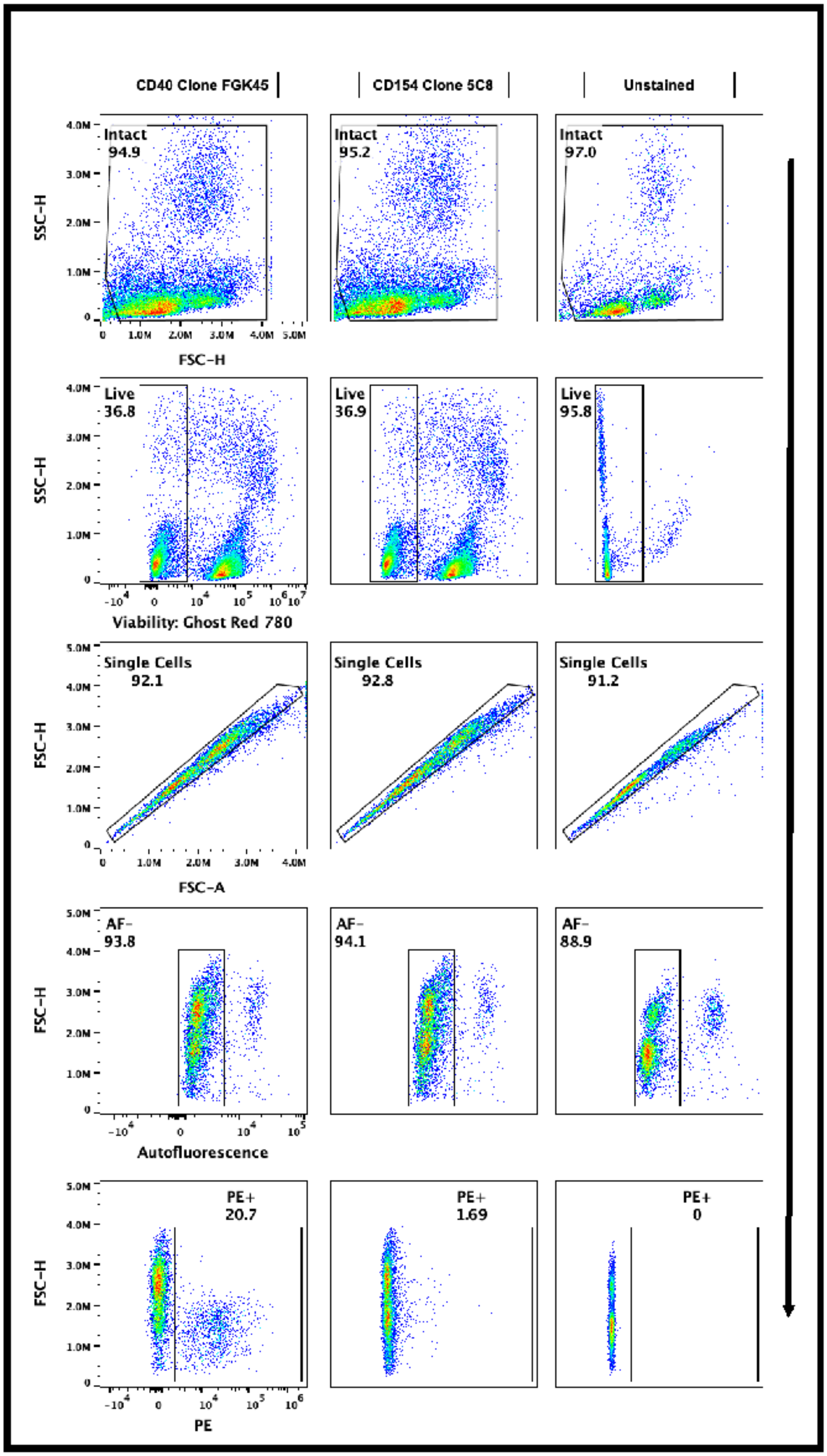
