## Supplemental Figure 5 for "Regulatory T Cell-like Response to SARS-CoV-2 in Jamaican Fruit Bats (*Artibeus jamaicensis*) Transduced with Human ACE2"

**Supplemental Figure 5**. Gating strategy to obtain CD4+ CD154+ splenocytes. Concatenated no peptide samples and FMOs (left) and concatenated peptide samples (right) were gated as follows: Intact Lymphocytes > Single Cells > Live Cells > CD4+ > CD154+. CVS files of CD4+CD154+ cells were then exported from FlowJo to obtain fluorescent intensities of CD154 on each cell for statistical analysis in Graph Pad Prism 9.

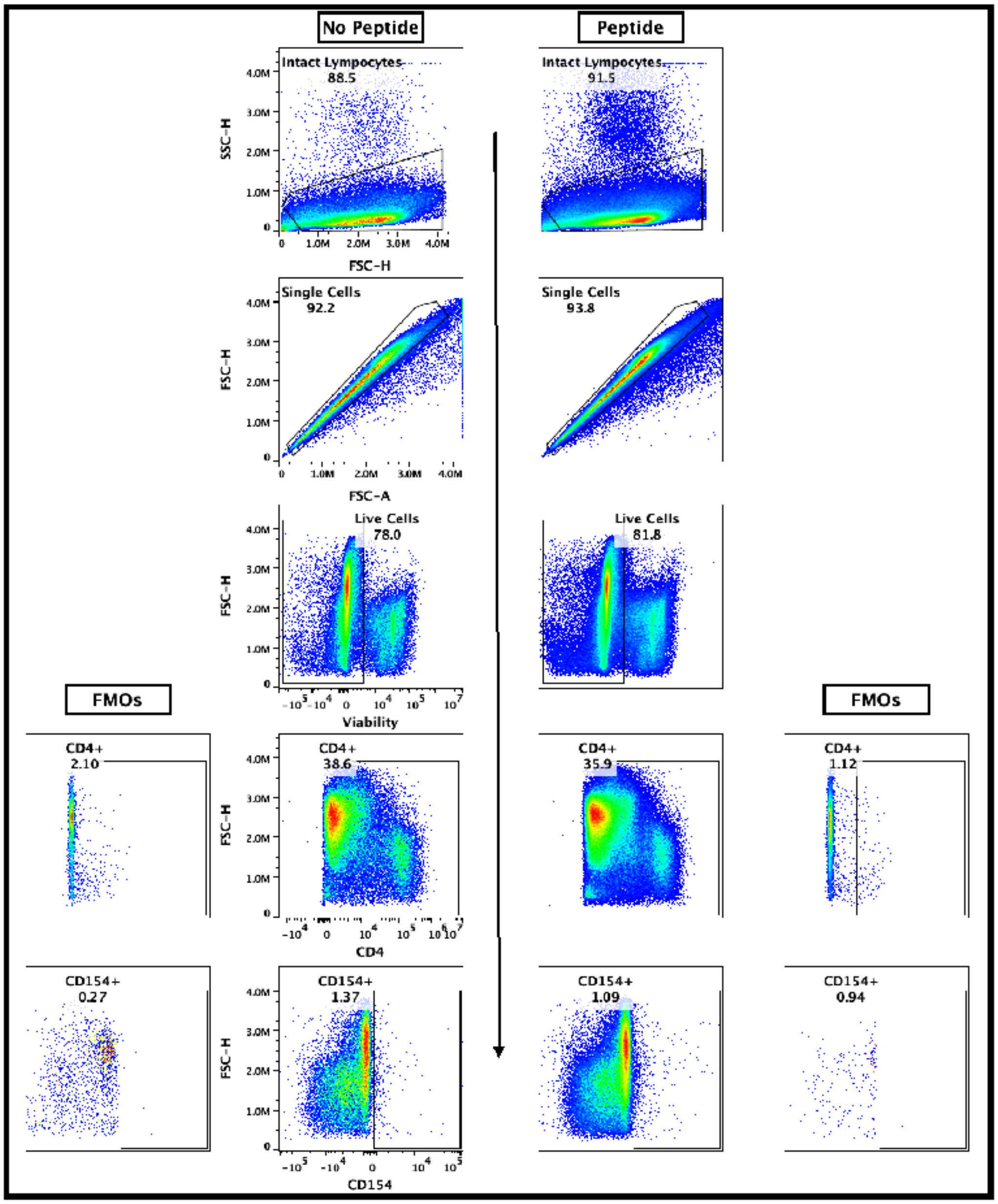
