## Supplemental Table 1 for "Regulatory T Cell-like Response to SARS-CoV-2 in Jamaican Fruit Bats (*Artibeus jamaicensis*) Transduced with Human ACE2"

**Supplemental Table 1.** Alignment of 20 ACE2 residues important for SARS-CoV-2 spike binding (Luan et al. and Wan et al. [cyan]). Residues identical to human are noted in black, whereas differences in red font. Species that are susceptible to SARS-CoV-2 are in red font, not susceptible in blue font, unknown susceptibility in black font.

| | ACE2 | AA position | | | | | | | | | | | | | | | | | | | | |
|---|---|---|---|---|---|---|---|---|---|---|---|---|---|---|---|---|---|---|---|---|---|---|
| **Common Name** | **Species name** | 24 | 27 | 28 | 30 | 31 | 34 | 35 | 37 | 38 | 41 | 42 | 45 | 82 | 83 | 330 | 353 | 354 | 355 | 357 | 393 | **ID** |
| Human | *Homo sapiens* | Q | T | F | D | K | H | E | E | D | Y | Q | L | M | Y | N | K | G | D | R | R | 20 |
| Syrian hamster | *Mesocricetus auratus* | Q | T | F | D | K | Q | E | E | D | Y | Q | L | N | Y | N | K | G | D | R | R | 18 |
| Deer mouse | *Peromyscus maniculatus* | Q | I | F | D | K | Q | E | E | D | Y | Q | L | N | Y | N | K | G | D | R | R | 17 |
| Intermediate horseshoe bat | *Rhinolophus affinis* | R | I | F | D | N | H | E | E | D | Y | Q | L | N | Y | N | K | G | D | R | R | 16 |
| Domestic cat | *Felis catus* | L | T | F | E | K | H | E | E | E | Y | Q | L | T | Y | N | K | G | D | R | R | 16 |
| Egyptian rousette | *Rousettus aegyptiacus* | L | T | F | E | K | T | E | E | D | Y | Q | L | T | Y | K | K | G | D | R | R | 15 |
| Pearson's horseshoe bat | *Rhinolophus pearsonii* | R | T | F | D | K | H | E | E | D | H | E | L | D | Y | N | K | D | D | R | R | 15 |
| Least horseshoe bat | *Rhinolophus pusillus* | L | K | F | N | D | S | E | E | D | Y | E | L | N | Y | N | K | G | D | R | R | 14 |
| American mink | *Neogale vison* | L | T | F | E | K | Y | E | E | E | Y | Q | L | T | Y | N | K | H | D | R | R | 14 |
| Ferret | *Mustela putorius* | L | T | F | E | K | Y | E | E | E | Y | Q | L | T | Y | N | K | R | D | R | R | 14 |
| Big-eared horseshoe bat | *Rhinolophus macrotis* | E | K | F | D | K | S | K | E | D | Y | E | L | N | Y | K | K | G | D | R | R | 13 |
| Chinese rufous horseshoe bat | *Rhinolophus sinicus* | E | I | F | D | K | T | K | E | D | H | Q | L | N | Y | N | K | G | D | R | R | 13 |
| Lander's horseshoe bat | *Rhinolophus landeri* | L | T | F | D | D | S | A | E | N | Y | Q | L | N | F | N | K | G | D | R | R | 13 |
| Jamaican fruit bat | *Artibeus jamaicensis* | D | T | F | E | K | T | E | E | E | Y | E | L | A | Y | N | K | N | D | R | R | 13 |
| House mouse | *Mus musculus* | N | T | F | N | N | Q | E | E | D | Y | Q | L | S | F | N | H | G | D | R | R | 13 |
| Greater horseshoe bat | *Rhinolophus ferrumequinum* | L | K | F | D | D | S | E | E | N | H | Q | L | N | F | N | K | G | D | R | R | 12 |
| Halcyon horseshoe bat | *Rhinolophus alcyone* | L | I | F | D | N | S | E | E | N | H | Q | L | K | F | N | K | N | D | R | R | 11 |
| Big brown bat | *Eptesicus fuscus* | E | I | F | Q | R | T | E | E | E | H | Q | L | R | Y | N | K | G | D | R | R | 11 |
| Brazilian free-tailed bat | *Tadarida brasiliensis* | E | I | F | Q | R | T | E | E | E | H | Q | L | R | Y | N | K | G | D | R | R | 11 |
| | | 24 | 27 | 28 | 30 | 31 | 34 | 35 | 37 | 38 | 41 | 42 | 45 | 82 | 83 | 330 | 353 | 354 | 355 | 357 | 393 | **ID** |
