## Supplemental Table 2 for "Regulatory T Cell-like Response to SARS-CoV-2 in Jamaican Fruit Bats (*Artibeus jamaicensis*) Transduced with Human ACE2"

**Supplemental Table 2.** Immune gene expression primers, 5' to 3'.

| Gene | Forward | Reverse |
|---|---|---|
| **Rps18** | GCGAGTACTCAACACCAATA | TTTTGGTGAGGTCGATGTCT |
| **IFNg** | CCTTGAAAGACAACCAGAGC | CCTCCATTTTGCTGTTGCTG |
| **IL-2** | CGAACTCAACCCTCTGAAGG | GTGCGTTCACTGTGACATTG |
| **IL-4** | AGCTCTGGTCTGCTTACTAG | TGGTCTCTTCTAAGGTGAGG |
| **IL-10** | TGAAAACAAGAGCAAAGCAG | TGAAAACAAGAGCAAAGCAG |
| **IL-12a** | TGCTGGCAGCTATTGATGAG | AGATCCGGTTCTTCAAGGGA |
| **IL-13** | CATCCAAAAGACCCAGAGG | ATCACTTCCACTTTGGTGTC |
| **IL-21** | AGATAATCAGCCTGCTAACG | CTCTGTTTCTGTCTTCTCCC |
| **IL-27** | GACCGTGAGTTTGGATCTCC | TCAGGGAGGTTGAATCCTGT |
| **TGFb** | TGTTCTTCAACACGTCGGAG | CGTGCTGTTCCACTTTCAAC |
| **Ebi3** | GCCCTACATGCTGAACATCA | TCTGGAGGGTCTGGTTTGAT |
| **IL-27Ra** | TGAGTCTTACCTGCCTTCCA | CTTGCTGGAATTGCTGTTGG |
| **gp130** | CTGAGTCCTTGAAGGCGTAC | TTCCCCACTTTCTTTGTCCG |
| **Tbx21** | CCAAAGGATTCCGGGAGAAC | AATTGACAGTTGGGTCCAGG |
| **Gata3** | ACCCCTGACTATGAAGAAGG | CACCTTTTTGCACTTTTTGG |
| **Foxp3** | CCAGCTCTGTGATCTGGAAC | AAAGCCTGTTGAGGTCCATC |
| **Cxcr4** | GCTCAGGCGACTATGACTCC | TGCCCACTATGCCAGTCAAG |
| **Cxcr5** | GGTCAGCCAACCTCATCACA | TAGAGGAAGCGGGAGGTGAA |
| **CD4** | CTCATTCCCACTCACCTTCG | CTCATTCCCACTCACCTTCG |
| **CD25** | TGTGAGTGCAAGAAAGGCTT | TTTTCCCAGAAAGAGTGGCC |
